# Elongator is required for root stem cell maintenance by regulating *SHORT ROOT* transcription

**DOI:** 10.1101/251603

**Authors:** Linlin Qi, Xiaoyue Zhang, Huawei Zhai, Jian Liu, Fangming Wu, Chuanyou Li, Qian Chen

## Abstract

*SHORTROOT* (SHR) is essential for stem cell maintenance and radial patterning in Arabidopsis thaliana roots, but how its expression is regulated is still unknown. Here, we report that Elongator regulates the transcription of *SHR*. The depletion of Elongator drastically reduced *SHR* expression and led to defective root stem cell maintenance and radial patterning. The importance of the nuclear localization of Elongator for its functioning, together with the insensitivity of the *elp1* mutant to the transcription elongation inhibitor 6-azauracil and the direct interaction of the ELP4 subunit with the C-terminal domain of RNA polymerase II (RNAPII CTD), support the notion that Elongator plays important roles in transcription elongation. Indeed, we found that ELP3 associates with the pre-mRNA of *SHR* and that mutation of Elongator reduces the enrichment of RNAPII on the *SHR* gene body. Moreover, Elongator interacted in vivo with SUPPRESSOR OF Ty4 (SPT4), a well-established transcription elongation factor that was recruited to the *SHR* locus. Together, these results demonstrate that Elongator acts in concert with SPT4 to regulate the transcription of *SHR*.

## Introduction

Root growth in higher plants relies on a group of pluripotent, mitotically active stem cells residing in the root apical meristem (RAM). In the RAM of the model plant *Arabidopsis thaliana*, the mitotically less active quiescent center (QC) cells, together with their surrounding stem cells, constitute the root stem cell niche (SCN), which continues to provide cells for all root tissues [1]. Pioneering studies have identified several key regulators that help determine the specification and functioning of the SCN. Among these, the GAI, RGA, and SCR (GRAS) family transcription factors SHORTROOT (SHR) and SCARECROW (SCR) provide positional information along the radial axis, whereas the plant hormone auxin, together with its downstream components, the PLETHORA (PLT) class of transcription factors, provides longitudinal information [2-5].

In addition to regulating the positional specification of the QC, SHR also controls the formative division of the cortex/endodermis initial (CEI) stem cell and its immediate daughter cell (CEID), which generates the separate endodermis and cortex cell layers constituting root ground tissue [1]. Interestingly, *SHR* is transcriptionally expressed in the stele, and its encoded protein moves into the outer adjacent cell layer, where its partner SCR sequesters SHR to the nucleus by forming the SHR-SCR complex [6, 7]. Recent efforts have successfully identified important transcriptional targets of the SHR-SCR complex. Among these are a group of so-called BIRD family genes encoding zinc finger proteins [8-10] and the cell-cycle gene *CYCLIND6;1* (*CYCD6;1*)[11].The spatiotemporal activation of *CYCD6;1* is controlled by a bistable switch involving SHR, SCR, and the cell differentiation factor RETINOBLASTOMA-RELATED (RBR) [12]. Despite these advances, how the master regulator gene *SHR* itself is regulated remains largely unknown.

In eukaryotic cells, protein-coding genes are transcribed by RNA polymerase II (RNAPII). The multifunctional protein complex, Elongator, was first identified as an interactor of hyperphosphorylated (elongating) RNAPII in yeast and was later purified from human and *Arabidopsis* cells [13-15]. Elongator consists of six subunits, designated ELP1 to ELP6, with ELP1 and ELP2 functioning as scaffolds for complex assembly, ELP3 acting as the catalytic subunit, and ELP4-6 forming a subcomplex important for substrate recognition [16-18]. In yeast, the loss of Elongator subunits leads to altered sensitivity to stresses including salt, caffeine, temperature, and DNA damaging agents [13, 19]. Since Elongator was copurified with elongating RNAPII and the ELP3 subunit showed histone acetylation activity, it was initially proposed that Elongator mainly functions as a transcription elongation factor, a process that occurs in the nucleus [13, 20, 21]. Shortly thereafter, this proposition was questioned, as several studies show that yeast Elongator has diverse functions related to its transfer (t)RNA modification activity thattake place in the cytoplasm [22-27].

The physiological functions of Elongator in mammals are exemplified by the finding that impaired Elongator activity in human is correlated with the neurological disorder familial dysautonomia [28] and that mutations in Elongator subunits are embryotic lethal in mice [29]. Like its yeast counterpart, human Elongator also has lysine acetyltransferase(KAT) activity. Among the major substrates for the KAT activity of human Elongator are Histone H3 and *α* -tubulin, reflecting the distinct functions of Elongator in the nucleus and cytoplasm. While, in the nucleus, acetylation of Histone H3 is linked to the function of Elongator in transcription [30], cytoplasmic acetylation of *α*-tubulin by Elongator underlies the migration and maturation of neurons [31].

Genetic studies have demonstrated that Elongator plays an important role in regulating multiple aspects of plant development and adaptive responses to biotic and abiotic stresses [15, 32-35]. Recent studies reveal the role of plant Elongator in regulating microRNA biogenesis and tRNA modification [36, 37].

Here, we report the action mechanism of plant Elongator in regulating root SCN and radial patterning. We show that the root developmental defects of Elongator mutants are largely related to drastically reduced *SHR* expression. We provide evidence that Elongator acts as a transcription regulator of *SHR*.

## Results

### Elongator is required for SCN maintenance and general root growth

To systematically evaluate the role of Elongator in regulating root growth, we investigated mutants of all six Elongator subunits (*elp1* to *elp6*, see Materials and Methods) and several double mutants including *elp1 elp2*, *elp1 elp4*, *elp1 elp6*, *elp2 elp4*, *elp2 elp6*, and *elp4 elp6*. Each of the single mutants exhibited similar reductions in root growth, and none of the investigated double mutant lines showed additive effects (Fig EV1A), implying that each subunit is essential for the functioning of Elongator and that Elongator acts as an integral complex that regulates root growth. Therefore, we used *elp1* as a representative mutant for detailed phenotypic analyses.

Cytological observations revealed that both cell division and cell elongation were reduced in *elp1* (Fig EV1B–H). In a Lugol’s iodine starch staining assay of WT roots expressing the QC-specific marker QC25, one layer of columella stem cells (CSCs) without starch staining was visible between the QC and the columella cell layers, hinting at a well-organized and functional SCN (Fig 1A). By contrast, in *elp1* root tips, QC25 expression was weak in the QC, but its expression pattern expanded downward and merged with that of starch staining, and the CSCs could not be clearly discerned (Fig 1B), suggesting the loss of QC cell identity and CSC differentiation. Consistently, in RNA *in situ* hybridization assays of WT roots, *WUSCHEL-RELATED HOMEOBOX 5 (WOX5)* was specifically expressed in the QC, but its expression pattern was diffuse and merged with neighboring cells in *elp1* roots (Fig 1C and D). These results, together with previous observations of the *elp2* mutant [35], indicate that Elongator is required for root SCN maintenance and general root growth.

**Figure 1.**
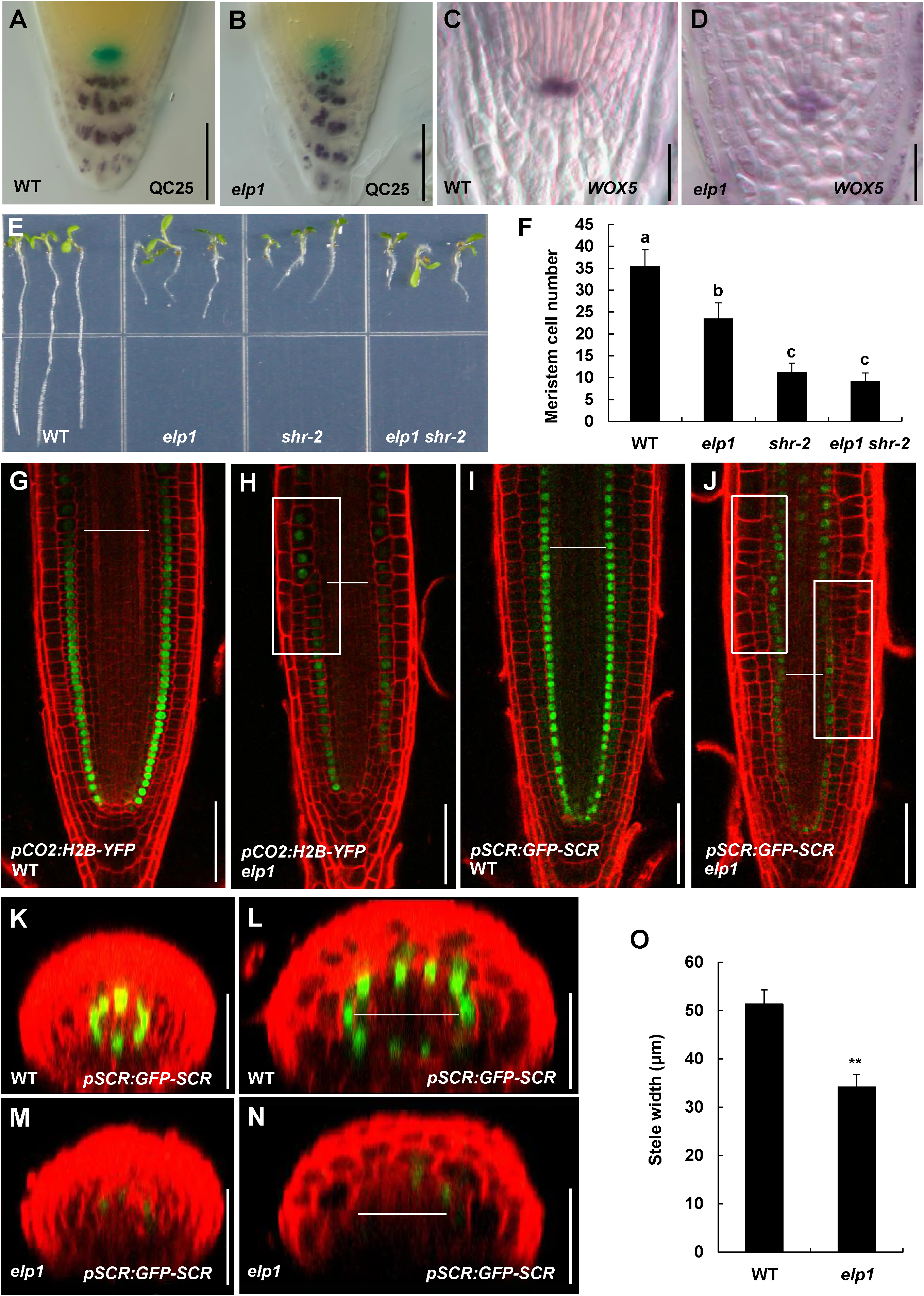
Elongator functions in root SCN maintenance and radial patterning through the SHR pathway. A, B Double staining of the QC-specific marker QC25 marker (blue) and starch granules (dark brown) in the wild type (WT) (A) and *elp1*(B)at 5 days after germination (DAG). C, D *WOX5* expression in 5-DAG WT (C) and *elp1*(D) revealed by whole-mount RNA *in situ* hybridization with a *WOX5* antisense probe. E Photographs of 5-DAGof WT, *elp1*, *shr-2*, and *elp1 shr-2*seedlings showing the involvement of the genetic relationship of Elongator with SHR in regulating root growth. F Quantification of meristem cell number of the indicated plants. Data shown are average and SD (*n* = 20). Samples with different letters are significantly different at *P*< 0.01. G, J Expression of the cortex-specific marker *pCO2:H2B-YFP* (G and H) and endodermis-specific marker *pSCR:GFP-SCR* (I and J) in WT (G and I) and *elp1*(H and J) roots. White rectangles highlight the disorganized cell layers in the *elp1* mutant, and horizontal white bars indicate the stele width (including pericycle cells). K, N Transverse confocal sections showing the expression of the endodermis-specific marker *pSCR:GFP-SCR* at the CEI/CEID position (K and M) and the transition zone (TZ) (L and N) in the WT (K and L) and *elp1* (M and N).Horizontal white bars indicate the stele width (including pericycle cells). O Quantification of stele width in WT and *elp1*. The stele width (including the pericycle cells) at the TZ position in the longitudinal confocal images was measured with ImageJ software. Data shown are average and SD (*n* = 20), and asterisks denote Student’s *t* test significance compared with the WT: \*\**P*< 0.01. Bars: A, B, and G–N, 50 μm; C and D, 20 μm.

### Elongator regulates radial patterning in roots through the SHR pathway

To investigate the genetic relationship between Elongator and the SHR pathway, we generated an *elp1 shr-2* double mutant line. At 5 DAG, general root growth and the meristem cell number of the double mutant were similar to those of *shr-2* (Fig 1E and F), indicating that *elp1* and *shr-2* do not have additive effects on root growth. These results support the notion that Elongator acts genetically in the SHR pathway to regulate root growth.

In parallel experiments, the *elp1 plt1-1 plt2-4* triple mutant line appeared to show an additive effect compared withits parental lines, *elp1* and *plt1-1 plt2-4* (Fig EV2A–F). Consistently, the *elp1* mutation had only a minor effect (if any) on *PLT1* and *PLT2* expression (Fig EV2G–J). These results support the notion that Elongator acts genetically in parallel with the PLT pathway.

In addition to having a defective SCN, *shr* mutants also exhibit irregular radial patterning and reduced stele width [8]. Hence, we examined whether *elp1* shows similar phenotypes. Using the cortex-specific marker *pCO2:H2B-YFP* and the endodermis-specific marker *pSCR:GFP-SCR*, we clearly distinguished these well-organized cell layers in WT roots (Fig 1G and I), whereas irregular patterning in certain regions of the cortex and/or endodermis cell layers was frequently observed in *elp1* roots (Fig 1H and J). Consistently, the expression levels of *CO2* and *SCR* were lower in *elp1* than in the WT (Fig 1G–N). Moreover, the stele width at the transition zone was also significantly reduced in *elp1* compared with the WT (Fig 1G–O). Together, the phenotypic similarity between *elp1* and *shr* strengthens the idea that Elongator acts in the SHR pathway to regulate radial patterning in roots.

### Depletion of Elongator impairs the expression of *SHR* and its target genes

Using the *SHR* promoter fusion line *pSHR:GFP* (for green fluorescent protein) and the SHR protein fusion line *pSHR:SHR-GFP*, we found that *SHR* expression levels were drastically reduced in *elp1* compared with the WT (Fig 2A–D). This observation was confirmed by RNA *in situ* hybridization (Fig 2E and F) and reverse-transcription quantitative polymerase chain reaction (RT-qPCR) assays (Fig 2G). Not surprisingly, as revealed by RT-qPCR, the expression levels of several SHR transcriptional targets, including *SCR*, *BR6ox2*, *CYCD6;1*, *MGP*, *NUTCRACKER* (*NUC*), *RECEPTOR-LIKE KINASE* (*RLK*), and *SNEEZY/SLEEPY 2* (*SNE*), were also substantially reduced in *elp1* compared with the WT (Fig 2G). We then investigated whether the *elp1* mutation impairs the expression of the cell-cycle gene *CYCD6;1*in the CEI/CEID. Previous studies have elegantly demonstrated that the spatiotemporal activation of *CYCD6;1* coincides with the formative division of CEI/CEID and that this process is strictly controlled by SHR and the related transcription factor SCR [11, 12]. As expected, the spatiotemporal expression of *CYCD6;1* in the CEI/CEID was largely disrupted in the *elp1* mutant (Fig 2H and I). Together, these results indicate that the depletion of Elongator impairs the expression of *SHR* and its target genes.

**Figure 2.**
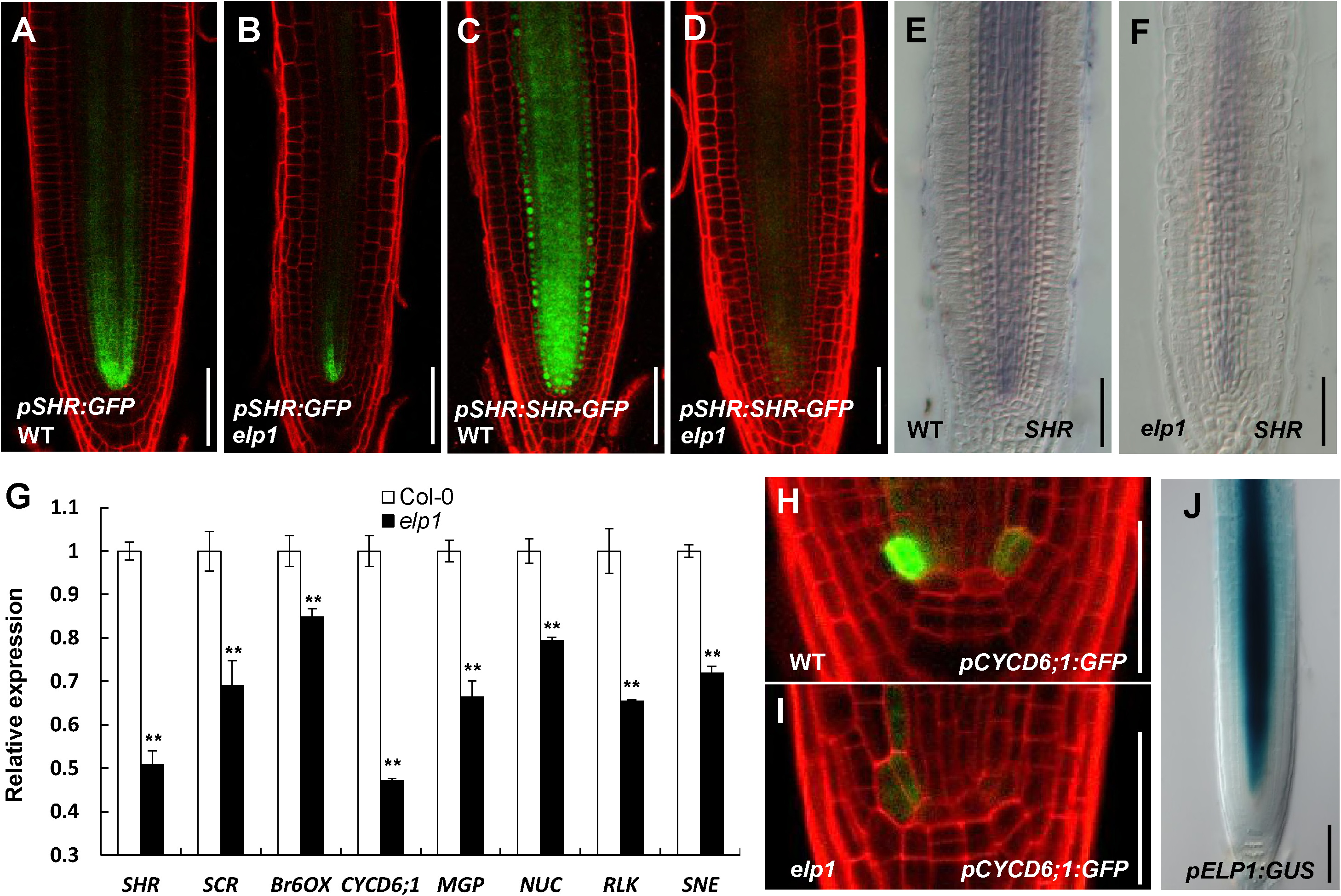
The effect of Elongator depletion on the expression of *SHR* and its target genes. A, F *SHR* expression is drastically reduced in the *elp1* mutant (B, D, and F) compared with the WT (A, C, and E), as revealed by the expression of the marker constructs *pSHR:GFP*(A and B) and *pSHR:SHR-GFP* (C and D) and by whole-mount RNA *in situ* hybridization with a *SHR* antisense probe (E and F). GRT-qPCR analysis showing the relative expression levels of *SHR* and its target genes in WT and *elp1*. Total RNA was extracted from 0.5cm root tip sections of 5-DAG seedlings. Transcript levels were normalized to the reference gene *PP2AA3*. Error bars represent SD (Student’s *t* test, \*\**P*< 0.01). The experiments were repeated three times, yielding similar results. H, I Representative images showing the location-specific expression and reduced expression of *pCYCD6;1:GFP* in CEI/CEID cells of the WT and the *elp1* mutant, respectively. JGUS staining of *pELP1:GUS* showing the expression pattern of *ELP1* in root tips. Data information: Bars in A–F, H, and I, 50 μm; J, 100 μm.

To visualize the expression pattern of the Elongator subunit ELP1, we fused the *ELP1*promoter with the -glucuronidase (GUS) reporter and generated *pELP1:GUS* transgenic plants. GUS staining revealed that, like*SHR*, *ELP1*washighly expressed in the stele of the root tip (Fig 2J). This observation strengthens the notion that Elongator regulates root development through the SHR pathway.

### Nuclear localization of Elongator is important for its function in regulating root development

To determine the subcellular localization of Elongator, we introduced the *pELP3:ELP3-GFP* fusion construct into *elp3*plants. The functionality of the ELP3-GFP fusion protein was verified by its ability to rescue the root growth defects of the *elp3* mutant (Fig EV3A). Confocal microscopy of *elp3;pELP3:ELP3-GFP* plants indicated that, in stele cells of the meristem region, ELP3-GFP was predominantly localized to the cytoplasm and, to a lesser extent, the nucleus (Fig 3A and B). Interestingly, we observed more obvious nuclear localization in columella cells and epidermis cells at the elongation zone in the previously reported line, *p35S:GFP-ELP3*[15](Fig EV3B).

**Figure 3.**
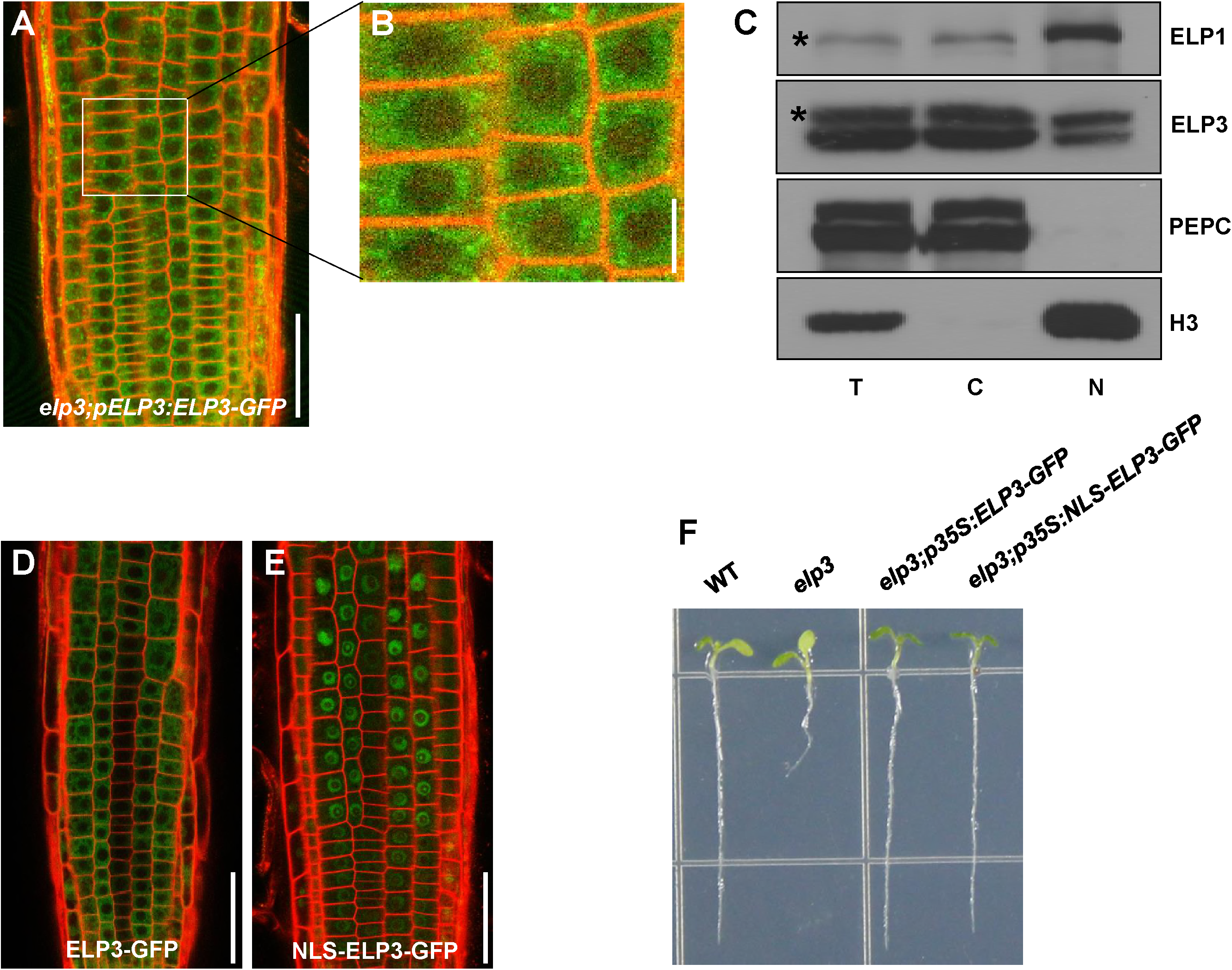
Importance of the nuclear localization of Elongator for its function. A, B Representative images showing the localization of ELP3-GFP in 5-DAG *elp3;pELP3:ELP3-GFP* transgenic plants. (B) Magnification of the image in the white rectangle in (A). C Cell fractionation assay of Col-0 seedlings. Ten-day-old Col-0 seedlings were collected for cell fractionation. Proteins from different fractions were immunoblotted with antibodies against ELP1, ELP3, H3, and PEPC. H3 and PEPC were used as nucleus- and cytoplasm-specific marker proteins, respectively. Asterisk indicates the position of the specific band. T, total extracts; C, cytoplasmic fraction; N, nuclear fraction. The experiments were repeated three times, yielding similar results. D, E Representative images showing the subcellular localization of ELP3-GFP (D) and NLS-ELP3-GFP (E), indicating the nuclear localization of NLS-ELP3-GFP. FPhotograph of 5-DAG seedlings showing that nuclear-localized ELP3 fully complemented the short-root phenotype of the *elp3* mutant. ELP3 was fused with an efficient SV40 nuclear localization signal (NLS) to artificially translocate it into the nucleus. The resulting constructs were transformed into the *elp3* background to determine phenotype complementation. Data information: Bars in A, E, and F, 50 μm; B, 10 μm.

We then employed a cell fractionation approach to determine the subcellular localization of endogenous ELP1 and ELP3. For these experiments, protein extracts of WT seedlings were fractionated and probed with antibodies that specifically recognize endogenous ELP1 or ELP3 protein (Fig EV4A and B). Histone H3 was exclusively detected in the nuclear compartment, whereas phosphoenolpyruvate carboxylase (PEPC) was only detected in the cytoplasmic compartment, validating our approach. Consistent with the above cytological observations, endogenous ELP1 and ELP3 were clearly detected in both the cytoplasmic and nuclear fractions (Fig 3C). We obtained similar results from cell fractionation experiments using *elp3;p35S:ELP3-myc* transgenic plants, in which the root growth defects of the *elp3* mutant had been rescued (Fig EV4C). These results help confirm the finding that *Arabidopsis* Elongator is located in both the cytoplasm and the nucleus.

To determine if the nuclear localization of ELP3 is critical for its function, we artificially confined ELP3 to the nuclear compartment and investigated whether it would still retain its function. For these experiments, *ELP3-GFP* was fused with the efficient nuclear localization sequence SV40 NLS [38] to generate the *p35S:NLS-ELP3-GFP* construct. Analysis of the resulting transgenic plants indicated that the NLS-ELP3-GFP fusion protein was successfully translocated into the nucleus and, more importantly, the nuclear-localized NLS-ELP3-GFP fusion protein was functional, as it fully rescued the root growth defects of *elp3* (Fig 3D–F). These results support the notion that the nuclear localization of Elongator is important for its function in regulating root development.

### Elongator functions as a transcription elongation factor to regulate *SHR* transcription

Our finding that the nuclear localization of plant Elongator is important for its biological function suggested that Elongator might act as a transcription elongation factor involved in RNAPII-dependent transcription. To investigate this notion, we first examined whether the Elongator subunits interact with the conserved C-terminal domain(CTD) of RNAPII, an interaction platform between RNAPII and other proteins involved in transcription.

RNAPII CTD interacted with ELP4 and ELP5 in yeast two-hybrid (Y2H) assays (Fig 4A). To determine whether ELP4 interacts with CTD *in planta*, we conducted firefly luciferase (LUC) complementation imaging (LCI) assays in *Nicotiana benthamiana* leaves. In these experiments, ELP4 was fused to the N-terminal half of LUC (nLUC) to produce ELP4-nLUC, whereas CTD was fused to the C-terminal half of LUC (cLUC) to produce cLUC-CTD. *N. benthamiana* cells co-expressing ELP4-nLUC and cLUC-CTD displayed strong fluorescence signals, whereas those co-expressing nLUC and cLUC-CTD or ELP4-nLUC and cLUC displayed no signal (Fig 4B and C), confirming that the ELP4–CTD interaction occurs *in vivo*.

**Figure 4.**
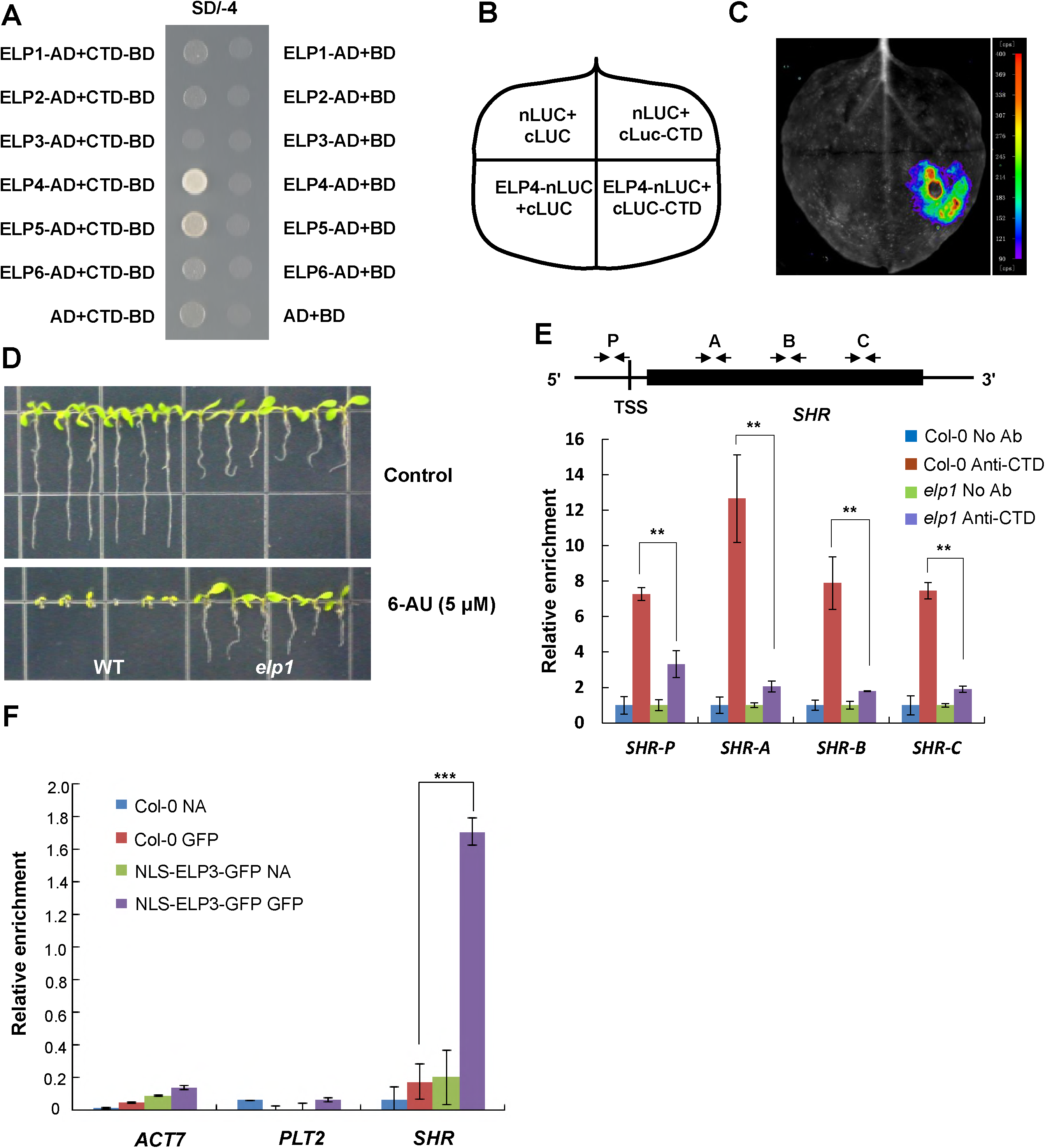
Elongator functions as a transcription elongation factor to regulate *SHR* expression. A Interactions of different Elongator subunits with RNAPII CTD in a yeast two-hybrid assay. The yeast transformants were dropped onto SD/-Ade/-His/-Trp/-Leu (SD/-4) medium to assess protein–protein interactions. The experiments were repeated three times with similar results. B, C Firefly luciferase complementation imaging assay showing the interaction of the ELP4 subunit with RNAPII CTD in *N. benthamiana*. *N. benthamiana* leaves were infiltrated with *Agrobacterium* containing the indicated construct pairs (B). The image was obtained 3 days after infiltration (C). The colored bar indicates the relative signal intensity. The experiments were repeated three times, yielding similar results. D Photographs of 5-DAG seedlings showing the insensitivity of the *elp1* mutant to 6-AU. WT and *elp1*seeds were sown on 1/2MS medium without or with 0.5 mg/L 6-AU, and the plates were photographed at 5 DAG. EChIP-qPCR results showing that the *elp1*mutation impairs the recruitment of RNAPII to various regions of the *SHR* locus. Chromatin was extracted from Col-0 and *elp1* seedlings at 5 DAG and precipitated with an anti-CTD antibody (Abcam). Precipitated DNA was amplified with primers corresponding to the different regions of *SHR* as shown. The “No Ab” (no antibody) precipitates served as negative controls. The ChIP signal was quantified as the percentage of total input DNA by qPCR and arbitrarily set to 1 in the “No Ab” samples. TSS indicates transcription start site. The experiments were repeated three times, yielding similar results. Error bars represent SD. Asterisks indicate significant differences between Col-0 and the *elp1* mutant, according to Student’s *t* test (\*\**P*< 0.01). FRIP-PCR results showing the association of Elongator with *SHR* mRNA, but not with *PLT2* mRNA. Protein-RNA complexes were isolated from Col-0 and *elp3;p35S:NLS-ELP3-GFP* seedlings at 5 DAG and precipitated with an anti-GFP antibody (Abcam). Precipitated RNA either with reverse transcription was amplified with primers targeting the respective CDS regions. The RIP signal was quantified as the percentage of total input RNA by qPCR. Samples before precipitation were taken as “Input”, and the “NA” (no antibody) precipitates served as negative controls. The experiments were repeated three times with similar results.

We then examined the response of *elp1* to 6-Azauracil (6-AU), an inhibitor of enzymes involvedin purine and pyrimidine biosynthesis. In yeast, 6-AUis a widely used inhibitor of transcription elongation, as it alters nucleotide pool levels *in vivo* [39]. Strikingly, *elp1* was more resistant to 6-AU-induced root growth inhibition than the WT (Fig 4D), providing another line of evidence that Elongator functions as a transcription elongation factor in *Arabidopsis*.

Next, we investigated whether Elongator participates in RNAPII-dependent transcription elongation of *SHR* using chromatin immunoprecipitation (ChIP)-qPCR assays. In WT plants, CTD was highly enriched on both the transcription start site (TSS) and gene body of *SHR* (Fig 4E). In the *elp1* mutant, however, CTD levels on the *SHR* locus were significantly reduced (Fig 4E), revealing that Elongator is important for the recruitment of RNAPII to the *SHR* locus during *SHR* transcription. This line also demonstrates that Elongator is mainly involved in transcription elongation, rather than trancription initiation.

We then performed RNA-immunoprecipitation (RNA-IP) assays to investigate whether Elongator associates with the pre-mRNA of *SHR*. Specifically, we used the GFP antibody to immunoprecipitate ELP3-GFP from extracts of the *elp3;p35S:NLS-ELP3-GFP* transgenic line. The resulting ELP3-GFP immunoprecipitates were then reverse transcribed into cDNAs and subjected to RT-PCR with primers specific for *SHR*, *PLT2*, or *ACTIN7* (*ACT7*). Pre-mRNA of *SHR* was detected in immunoprecipitates from the *elp3;p35S:NLS-ELP3-GFP* line but not from those of the WT (Fig 4F), confirming that ELP3 indeed associates with *SHR* pre-mRNA. As a control, *PLT2* and *ACT7*pre-RNAs were not detected in the same ELP3-GFP immunoprecipitates (Fig 4F), suggesting the ELP3 specifically associates with the pre-mRNA of *SHR*. Together, these results led us to conclude that Elongator regulates the transcription of *SHR* through associating with its pre-mRNA.

### ELP1 associates with the transcription elongation factor SPT4, which is recruited to the *SHR* locus

Our results support the notion that Elongator regulates the transcription elongation of *SHR* through associating with the pre-mRNA of *SHR*. Intriguingly, however, we failed to detect Elongator enrichment on the *SHR* locus in the ChIP experiments. We speculated that Elongator might act in concert with other known transcription elongation factors to regulate the elongation of *SHR* transcript. Indeed, ELP1 and ELP3 were recently affinity copurified with SUPPRESSOR OF Ty4 (SPT4)and several other conserved transcription elongation factors in eukaryotic cells [40, 41]. In *Arabidopsis*, SPT4 is encoded by two redundant genes, designated *SPT4-1* and *SPT4-2* [40]. In a coimmunoprecipitation (Co-IP) assay using *p35S:SPT4-2-GFP* plants and anti-ELP1 antibodies, SPT4-2-GFP pulled down native ELP1, indicating that ELP1 associates with SPT4-2 *in vivo* (Fig 5A). The *in vivo* association of ELP1 with SPT4-2 was further confirmed in LCI assays: *N. benthamiana* cells co-expressing ELP1-nLUC and cLUC-SPT4-2 displayed strong fluorescence signals (Fig 5B and C).

**Figure 5.**
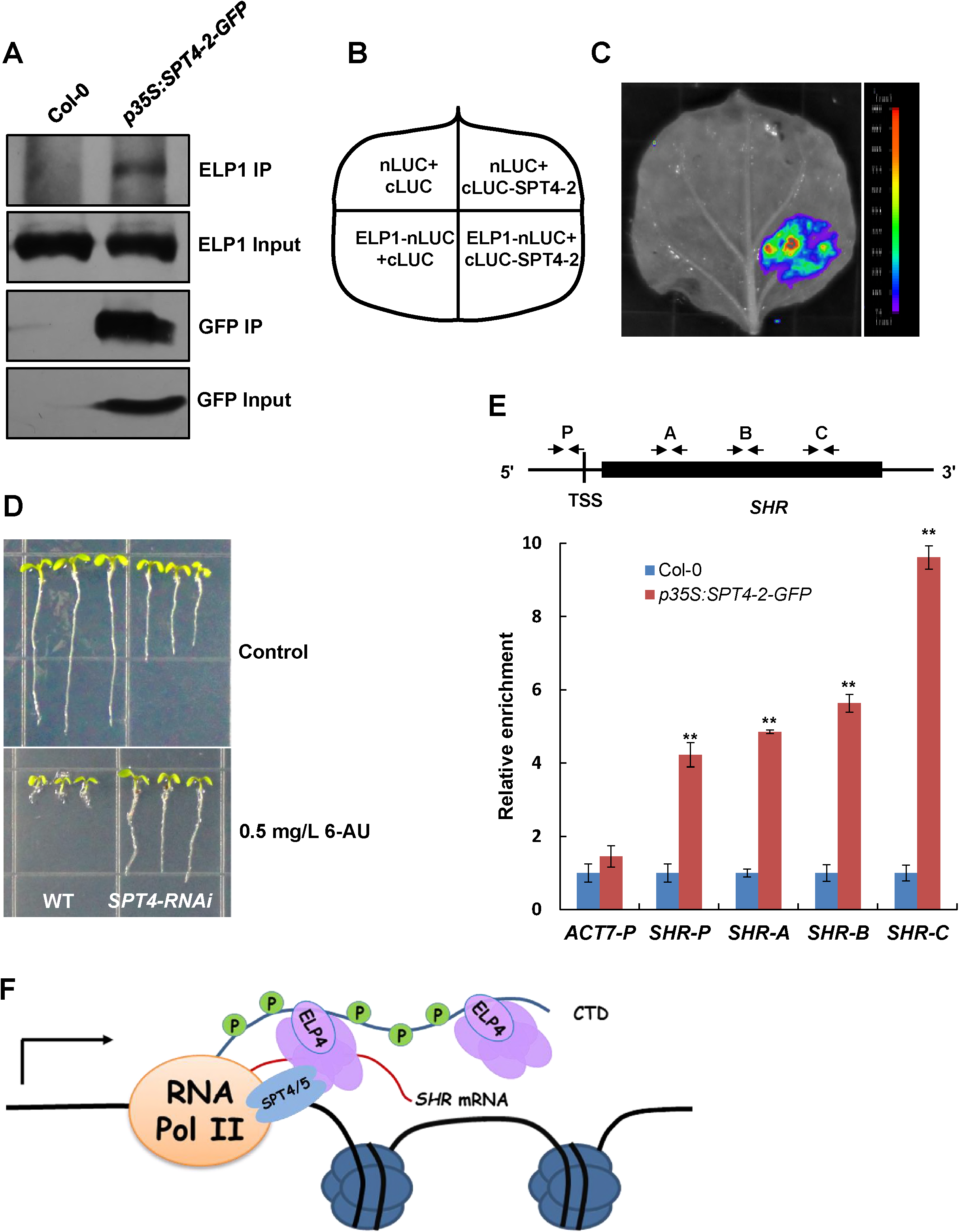
Elongator functions in concert with SPT4/SPT5. ACo-IP assay showing that Elongator associates with SPT4-2 in plant cells. Protein extracts from 10-day-old Col-0 and *p35S:SPT4-2-GFP* seedlings were immunoprecipitated with an anti-GFP antibody (Abcam). Samples before (Input) and after immunoprecipitation (IP) were blotted with anti-GFP and anti-ELP1 antibodies. The experiments were repeated three times with similar results. B, C Firefly luciferase complementation imaging assay showing the interaction of the ELP1 subunit with SPT4-2 in *N. benthamiana*. *N. benthamiana* leaves were infiltrated with *Agrobacterium* strains containing the indicated construct pairs (B). The image was obtained 3 days after infiltration (C). The colored bar indicates the relative signal intensity. The experiments were repeated three times, yielding similar results. D Photographs of 5-DAG seedlings showing the insensitivity of *SPT4-RNAi* to 6-AU. WT and *SPT4-RNAi*seeds were sown on 1/2MS medium without or with 0.5 mg/L 6-AU, and the plates were photographed at 5 DAG. EChIP-qPCR results showing the enrichment of SPT4-2 on the *SHR* locus. Sonicated chromatin from 5-DAG Col-0 and *p35S:SPT4-2*-GFP seedlings was precipitated with an anti-GFP antibody (Abcam). The precipitated DNA was used as template for qPCR analysis with primers targeting different regions of the *SHR* locus as shown. The promoter region of *ACT7* (*ACT7-P*) was used as a negative control. The ChIP signal was quantified as the percentage of the total input DNA and was arbitrarily set to 1 in Col-0. TSS, transcription start site. The experiments were repeated three times, yielding similar results. Error bars represent SD. Asterisks indicate significant differences, according to Student’s *t* test, \*\**P*< 0.01. F Proposed mechanism in which Elongator acts in concert with SPT4/SPT5 to regulate the transcription elongation of *SHR*. During the transcription elongation process of *SHR*, Elongator directly interacts with the RNAPII CTD and the nascent *SHR* mRNA and associates with SPT4/SPT5. Meanwhile, SPT4/SPT5 is in close contact with the chromosomal region harboring the *SHR* locus, the RNAPII subunits, and Elongator.

To demonstrate that SPT4-2 plays a role in root development, we generated *SPT4-RNAi* plants in which the expression of both *SPT4-2* and *SPT4-1* was knocked down by RNA interference (RNAi) (Fig EV5). Like*elp1* and the other *elp* mutants (Fig EV1), the *SPT4-RNAi*plantsalso showed reduced root growth (Fig 5D). These results are consistent with a previous observation, and they suggest that the interplay between Elongator and SPT4/SPT5 might help regulate root development [18, 40-42]. Furthermore, like the*elp1* mutant, *SPT4-RNAi* plants were less sensitive to 6-AU-induced root growth inhibition than WT plants (Fig 5D), implying that the function of SPT4-2 in regulating root growth is related to transcription elongation. Indeed, the ChIP-qPCR assays revealed that SPT4-2 was enriched on the *SHR* locus and importantly, the levels of SPT4-2 on the *SHR* gene body regions were higher than those on the *SHR* promoter region (Fig 5E), suggesting that SPT4 is mainly involved in *SHR* transcription elongation. Taken together, our results support the notion that Elongator acts in concert with SPT4/SPT5 to regulate the transcription of *SHR*, thereby regulating root development.

## Discussion

### Elongator is required for root SCN maintenance and radial patterning through regulating *SHR* gene expression

In addition to reduced primary root growth and defective SCN reported previously [35], we observed irregular radial patterning and reduced stele width in Elongator single and double mutants, indicating that Elongator functions as an integral complex in the regulation of root development. Several lines of evidence indicate that the developmental defects in roots of the Elongator mutants are largely due to an impaired SHR pathway. Indeed, the *elp1* root phenotypes resemble those of *shr* (Fig 1) and coincide with the similar developmental gene expression patterns in the primary root, with the highest expression in the stele tissue of root tips. Elongator acts upstream of SHR and independently of the PLT pathway, as the expression levels of *SHR* and its target genes *SCR* and *CYCD6;1* were drastically reduced in *elp1* (Fig 2A–I), whereas the expression of *PLT1* and *PLT2* was less affected in this mutant (Fig EV2). Thus, Elongator regulates root development mainly through its impact on the SHR pathway. In contrast to our knowledge about SHR target genes and interacting proteins, little had been known about how *SHR* itself is regulated, except that *SHR* expression is not regulated by PLT1, PTL2, SCR, or SHR itself [3, 4]. Our results indicate that Elongator is a regulator of *SHR* expression. The developmental and external stimuli that regulate the expression of the genes encoding the six Elongator subunits, such as hormones and temperature [18], might affect the production of Elongator subunits, thus affecting the assembly and accumulation of this crucial complex and thereby controlling (to some extent) *SHR* gene expression. Elongator might serve as an interface between these stimuli and SHR, and the regulation of the *SHR* gene at the transcription elongation stage might render its expression more flexible and responsive, which is important for the spatial and temporal control of root development.

### Elongator regulates transcription of *SHR* in concert with SPT4/SPT5

The role of Elongator as a transcription elongation factor is highly controversial in yeast and human[17] and was recently suggested in plants[18]. Here, using various experimental approaches, we provide direct evidence for the transcription elongation activity of Elongator. Using confocal microscopy analysis of various transgenic lines combined with cell fractionation, we showed that the Elongator subunits are partially localized to the nucleus, although major portions of these subunits are localized to the cytoplasm (Figs 3A–C, 3B and 4C). Moreover, the nuclear localization of these subunits is important for their biological function, as artificially nucleus-localized ELP3 was still able to complement the *elp3* mutant phenotype (Fig3D–F); a similar experiment was performed in yeast, but with contrasting results [43]. Our protein purification assays revealed a direct interaction of the ELP4 subunit with RNAPII CTD (Fig 4A–C) and the association of ELP1 with the well-established transcription elongation factor SPT4/SPT5 (Fig 5A–C), indicating that Elongator is involved in transcription elongation. However, we failed to detect an interaction between ELP4 and RNAPII in a Co-IP assay. This result confirms a previous report showing interactions between the different subunits of the Elongator but no interactions between Elongator subunits and RNAPII using TAP-MS (tandem affinity purification/mass spectroscopy) [15], but disagrees with a more recent report showing co-purification of the two largest subunits of RNAPII with Elongator subunits ELP1 and ELP3 using SPT4 as bait [40]. Hence, the association of Elongator with RNAPII *in vivo*might be transient and dynamic [42].

Yeast mutants defective in transcription elongation exhibit an altered sensitivity to 6-AU [44, 45]. We found that both *elp1* and *SPT4-RNAi* plants were highly resistant to 6-AU treatment (Figs 4D and 5D), suggesting that Elongator plays a similar role in transcription elongation to that of SPT4/SPT5. Finally, we detected a significant reduction in the enrichment of RNAPII CTD on the *SHR* locus in *elp1* compared with the WT (Fig 4E), as well as an association of ELP3 with *SHR* mRNA (Fig 4F). Moreover, its associated protein, SPT4, was also recruited to *SHR* chromatin (Fig 5E). These findings indicate that both Elongator and SPT4/SPT5 are directly involved in the transcription regulation of *SHR*. However, we failed to detect significant enrichment of Elongator on the *SHR* region via ChIP, possibly due to a lack of DNA-binding activity and/or the dynamic properties of its interaction with other proteins. No ChIP data are currently available for the enrichment of Elongator on transcriptional regulatory regions in *Arabidopsis*. Hence, we propose a model in which Elongator acts in concert with SPT4/SPT5 to maintain the transcription of *SHR*. In this model, Elongator directly interacts with the RNAPII CTD through the ELP4 subunit and associates with *SHR* mRNA, whereas SPT4/SPT5 associates with RNAPII and Elongator, as well as the chromosomal region harboring *SHR* (Fig 5F).

Transcription elongation is a tightly controlled, dynamic process that can be divided into three distinct stages: promoter escape, promoter-proximal pausing, and productive elongation. Based on their activities, transcription elongation factors can be categorized as positive or negative. In yeast, mutations of positive transcription elongation factors are often associated with hypersensitivity to 6-AU, whereas the disruption of negative transcription elongation factors renders the cells less sensitive to 6-AU [45]. We demonstrated that Elongator associates with SPT4/SPT5 (Fig 5A–C) and that both *elp1* and *SPT4-RNAi* are highly insensitive to 6-AU (Figs 4D and 5D), suggesting that Elongator and SPT4/SPT5 arenegative transcription elongation factor, as in yeast. Indeed, human SPT4/SPT5, i.e., 5,6-dichloro-1-β-D-ribofuranosylbenzimidazole(DRB) sensitivity-inducing factor (DSIF),is a negative transcription elongation factor that contributes to promoter-proximal pausing [46]. Transcriptional pausing is also thought to participate in mRNA synthesis, possibly through the formation of a “preactivated” state [46]. Hence, Elongator and SPT4/SPT5 might play an (as yet) unknown role in promoter-proximal pausing in *Arabidopsis*.

### Functional diversification of Elongator in eukaryotes

In yeast, tRNA modification appears to be the direct, unequivocal biochemical function of Elongator, because its cytoplasmic localization [43] excludes the possibility of this complex being a transcription elongation factor. In addition, overexpressing two related tRNA species rescued almost all of the reported phenotypes for yeast Elongator mutants, including reduced H3K14Ac levels, suggesting that even histone acetylation might be an indirect effect of Elongator activity [23]. Various biological processes are modulated through Elongator tRNA modifications, such as telomeric gene silencing [25], cell-cycle control [26], and oxidative stress responses [27]. However, in animal cells, Elongator is partially localized to the nucleus [14, 31], and its role in transcription elongation is supported by the enrichment of Elongator subunits on certain genes related to cell migration. Here, we provided direct evidence that Elongator acts as a transcription regulator in *Arabidopsis*. Since Elongator was first copurified with elongating RNAPII from total cell extracts in yeast [13], and their interaction was further confirmed in human cells [14] and here in *Arabidopsis*, it is reasonable to suspect that the association of Elongator with RNAPII in yeast is not merely a coincidence. Thus, we propose that both tRNA modification and RNAPII association are two functions of Elongator. However, the cytoplasmic localization of this complex in yeast precludes it from being a transcriptional regulator, and therefore the tRNA modification activity of Elongator contributes the most to its function. By contrast, in human and plant cells, the acquisition of a partial nuclear localization for this complex might have caused it to develop a capacity for transcriptional regulation. In animals, the cytoplasm-localized Elongator evolved various other activities, such as acetylation of α-tubulin in the mouse cortex [31]. In plants, Elongator also plays roles in the cytoplasm, namely, its tRNA modification activity is conserved in *Arabidopsis*[16], but how this localization contributes to its biological function is still uncertain, although auxin responses depend on Elongator tRNA activity [37].

Transcriptional regulation is a major, delicate regulatory mechanism involving numerous proteins. The key players in postembryonic root development in plants, such as PLT1, PLT2, SHR, SCR, and WOX5, are all transcription factors [5]. Several other transcriptional regulators are also required for this process, such as chromatin-remodeling factors [5], histone acetyltransferase, and splicing factors. In plants, Elongator might play a role in the crosstalk between environmental and developmental stimuli to flexibly control *SHR* transcription, thereby modulating root growth and development throughout the plant’s lifecycle.

## Materials and Methods

### Plant Material and Growth Conditions

The Elongator mutants *elp1* (*abo1-2*, SALK_004690), *elp2*, *elp4* (SALK_079193), *elp6*and the double mutants *elp1 elp2*, *elp1 elp4*, *elp1 elp6*, *elp2 elp4*, *elp2 elp6*, and *elp4 elp6*(Zhou et al., 2009) were obtained from Zhizhong Gong Lab. The mutants *elp3*(*elo3-6*, GABI_555H06) (Nelissen et al., 2010) and *elp5* (GABI_700A12) were ordered from the Arabidopsis Biological Research Center. The following mutants and marker lines *plt1-4 plt2-2*, *shr-2*, *pCYCB1;1:GUS*, QC25, *pSHR:GFP*, *pSHR:SHR-GFP*, *pSCR:GFP-SCR*, *pCO2:H2B-YFP*, *pCYCD6;1:GFP* were kindly provided by Ben Scheres and maintained in our lab for years. The triple mutant *elp1 plt1-4 plt2-2*, the double mutant *elp1 shr-2*, and different marker lines in the mutant background were all obtained by genetic crossing.

Seeds of *Arabidopsis thaliana* (L.) were surface-sterilized with 10% (v/v) bleach for 10 min and washed three times with sterile water. Sterilized seeds were suspended in 0.1% (w/v) agarose and plated on half-strength Murashige and Skoog (1/2MS) medium. After stratification for 2 days at 4°C, they were transferred to the growth chamber at 22°C with a 16-h light/8-h dark cycle.

### Plasmid Construction and Plant Transformation

Approximately a 2.0-kb fragment including the promoter region and the coding sequences for the N-terminal 29 amino acids of *ELP1* was amplified from the genomic DNA by PCR and cloned into the *Pac*I*/Asc*I sites of the binary vector *pMDC162*, resulting in the *pELP1:GUS* construct, in which the coding sequences were fused in frame with GUS. For the *pELP3:ELP3-GFP* construction, the region containing the GFP-coding sequence and NOS-T fragment from the *pGFP-2* vector, the CDS of *ELP3* and its promoter sequences were sequentially cloned in frame into the binary vector *pCAMBIA1300* with the restriction enzyme cloning method. For generation of the *p35S:ELP3-myc* or *p35S:ELP3-GFP* plasmids, the *ELP3* CDS was first cloned with the pENTR Directional TOPO Cloning Kit (Invitrogen) and then recombined with the binary vector *pGWB17* or *pGWB5* respectively with the Gateway LR Clonase Enzyme Mix (Invitrogen). The *p35S:NLS-ELP3-GFP* construct was generated same as *p35S:ELP3-GFP* except that the sequence encoding the functional SV40 NLS[38] was attached to the forward primer used for cloning the ELP3 CDS. Construction method for the *p35S:SPT4-2-GFP* plasmid was the same as that for *p35S:ELP3-GFP*. For the *SPT4-RNAi* construct, the *SPT4-2* CDS was cloned into *pHellsgate 2* in both forward and reverse directions with a one-step BP reaction. All the primers used for the molecular cloning are listed in S1 Table.

All the constructs were transformed into the *Agrobacterium tumefaciens* strain GV3101 (pMP90) that was used for plant transformation with the vacuum infiltration method.

### Histology and Microscopy

Phenotypic analysis, Lugol staining, GUS staining, microscopic observation, and confocal microscopy were all done as described previously[47]. For marker expression control, at least 15 seedlings were used for each sample and representative images were shown. For quantitative measurements, 20 seedlings of each sample were analyzed and the statistical significance was evaluated by the Student’s *t* test. For multiple comparisons, an analysis of variance was followed by Fisher’s least significant difference test (SPSS) on the data.

### Whole-Mount RNA *in Situ* Hybridization

Whole-mount RNA was hybridized *in situ* according to the method previously described, and the probes for *WOX5*, *PLT1*, and *PLT2* had already been synthesized[47]. The antisense and sense probes for *SHR* were synthesized with digoxigenin-11-UTP (Roche Diagnostics) by T7 RNA polymerase from an *SHR*-specific fragment with the T7 promoter sequence either at the reverse primer or at the forward primer, respectively (Appendix Table 1). To enhance the probe permeability for the *SHR* detection, *SHR* probes were hydrolyzed to an approximately 100-bp size, the proteinase K concentration was increased to 80 μg/mL, and the incubation time was prolonged to 20 min.

### Gene Expression Analysis

For RT-qPCR analysis, approximately 0.5-cm root tips were harvested from 5-day-old seedlings for RNA extraction with TRIzol reagent (Invitrogen). First-strand cDNA was synthesized from 2 μg of total RNA with the M-MLV reverse transcriptase (Promega) and oligo(dT) primer and was quantified with the LightCycler 480 II apparatus (Roche) and the SYBR Green Kit (Takara) according to the manufacturer’s instructions. The expression levels of the target genes were normalized to the reference gene *PP2AA3*. The statistical significance was evaluated by Student’s *t* test. Primers used for RT-qPCR analysis are listed in Appendix Table 1, of which some had been described previously[11].

### Cell Fractionation

Cell fractionation was performed according to the CELLYTPN1 protocol (Sigma) with slight modifications. Briefly, 4 g of 10-day-old WT seedlings was harvested and fully ground in liquid nitrogen. The powder was transferred to 8 mL of precooled NIBA buffer and filtered through nylon membrane. Triton X-100 was added to a final concentration of 0.5% (v/v) and the sample was kept on ice for 15 min, followed by centrifugation at 2,000×g at 4°C for 10 min. The extracts before centrifugation were collected as total proteins (T), whereas the supernatants after centrifugation were collected as cytoplasmic fractions (C). The pellets were resuspended in 1 mL of NIBA and applied on top of a 800-μL cushion of 1.5 M sucrose, followed by centrifugation at 12,000×g at 4°C for 10 min. The pellets were washed twice by resuspension in 1 mL of NIBA and centrifugation at 12,000×g for 5 min. Hereafter, the pellets were resuspended in 600 μL of NIBA as nuclear fraction (N). For each fraction, samples of 20 μL of protein were used for Western blot analysis.

### Antibody Preparation

The partial CDS encoding the 400 amino acids of the C-terminus of ELP1 and the full-length CDS of ELP3 were cloned into the *pET-28a* vector to express the recombinant proteins in *Escherichia coli* strain BL21. The primers used for cloning are listed in S1 Table. The recombinant proteins were used to raise polyclonal antibodies in mouse.

### Western Blot Analysis

Protein extraction and Western blot were done according to standard protocols. Seedlings were ground into a fine powder in liquid nitrogen and then transferred to extraction buffer (50 mM Tris-HCl, pH 8.0, 150 mM NaCl, 1% [v/v] Nonidet P-40, 1 mM phenylmethylsulfonyl fluoride [PMSF], 10 μM MG132, and protease inhibitor cocktail [Roche]). For Western blot analysis, protein samples were boiled for 5 min after mixing with sodium dodecyl sulfate (SDS) loading buffer, separated by SDS-polyacrylamide gel electrophoresis, and transferred to polyvinylidene fluoride membranes. Immunoblots were probed with the following antibodies: α-CTD (Abcam, 1:2000); α-H3 (Abcam, 1:4000); α-PEPC (Rockland, 1:2000); α-myc (Abmart, 1:2000); α-ELP1 (1:2000); and α-ELP3 (1:4000). Ponceau S-stained membranes were shown as loading controls.

### Y2H Assays

Y2H assays were based on the MATCHMAKER GAL4 Two-Hybrid System (Clontech). The full-length CDS of the six Elongator subunits was cloned into *pGADT7*, whereas the sequence encoding the RNAPII CTD was cloned into *pGBKT7*. The primers used for cloning are listed in Supplemental Table 1. Constructs were cotransformed into the yeast strain *Saccharomyces cerevisiae* AH109. The presence of the transgenes was confirmed by growth on SD/-Leu/-Trp plates. For protein interaction assessment, the transformed yeast was suspended in liquid SD/-Leu/-Trp medium and cultured to an optical density (OD) of 1.0. Five microliters of suspended yeast were dropped on plates containing SD/-Ade/-His/-Leu/-Trp medium. Interactions were observed after 3 days of incubation at 30°C.

### LCI Assays

LCI assays were done with *Nicotiana benthamiana* leaves as previously described[48]. Briefly, the full-length CDS of the two proteins were cloned into *pCAMBIA1300-NLUC* and *pCAMBIA1300-CLUC*. Primers used for the vector construction are shown in Supplemental Table 1. The resulting constructs were introduced into *Agrobacterium* strain GV3101. The tobacco leaves were coinfiltrated with combinations of strains as described and incubated for 3 days before observation with NightOWL II LB 983 (Berthold) imaging system.

### Co-IP Assays

The Co-IP assays were done according to the published method[49] with minor modifications. Briefly, total proteins were extracted from 10-day-old Col-0 and *p35S:SPT4-2-GFP* seedlings with the protein lysis buffer (50 mM Tris-HCl, pH 7.5, 150 mM NaCl, 0.1% [v/v] Triton X-100, 0.2% [v/v] Nonidet P-40, 0.6 mM PMSF, 20 μM MG132, and protease inhibitor cocktail [Roche]). To preclear 2-mg protein extracts, 20 μL protein A/G plus agarose (Santa Cruz) was used. Hereafter, the supernatants were incubated with 2 μL GFP antibody (Abcam) overnight and further precipitated with another 20 μL protein A/G plus agarose (Santa Cruz). The precipitated samples were washed four times with the lysis buffer and then eluted by boiling for 5 min with SDS loading buffer. Immunoblots were detected with α-ELP1 (1:2000) and α-GFP (Abmart, 1:2000).

### ChIP-qPCR Assays

ChIP assays were done according to the published protocol[49]. Briefly, 2 g of 5-day-old seedlings were cross-linked in 1% (v/v) formaldehyde for chromatin isolation. For immunobinding, 2 μL CTD antibody (Abcam) or GFP antibody (Abcam) was used. The protein-DNA complex was captured with 50 μL protein A agarose/salmon sperm DNA (Millipore). The eluted DNA was purified with QIAquick PCR purification kit (Qiagen) and used for qPCR analysis. Primers used for ChIP-qPCR are listed in Supplemental Table 1.

### RIP-PCR Assays

The RIP assays described previously[50] were slightly modified. Five-day-old seedlings of *elp3;p35S:NLS-ELP3-GFP* were harvested for RIP assays. The seedlings were cross-linked in 1% (v/v) formaldehyde. Subsequently, protein-RNA complexes were isolated and immunoprecipitated according to published procedures. The associated RNAs were detected with semi-quantitative reverse transcription PCR with primer pairs listed in S1 Table.

### Accession Numbers

The sequence data can be found in the *Arabidopsis* Genome Initiative under the following accession numbers: *ELP1* (At5g13680), *ELP2* (At1g49540), *ELP3* (At5g50320), *ELP4* (At3g11220), *ELP5* (At2g18410), *ELP6* (At4g10090), *SHR* (AT4g37650), *SCR* (AT3g54220), *BR6ox2* (AT3g30180), *CYCD6;1* (At4g03270), *MGP* (AT1g03840), *NUC* (AT5g44160), *RLK* (At5g67280), *SNE* ((At5g48170), *PP2AA3* (AT1g13320), *PLT1* (At3g20840), *PLT2* (At1g51190), *WOX5* (At3g11260), *ACT7* (At5g09810), *SPT4-1* (At5g08565), and *SPT4-2* (At5g63670).

## Acknowledgments

We thank Zhizhong Gong, Ben Scheres, and Klaus Palme for sharing their research materials and Martine De Cock for helpingus prepare the manuscript. This work was supported by the Tai-Shan Scholar Program from the Shandong Province, the State Key Laboratory of Plant Genomics of China, and the State Key Lab of Crop Biology; the National Basic Research Program of China (2013CB967301 2015CB942900); the National Natural Science Foundation of China (31320103910).

## Author contributions

**Conceptualization:** CL LQ QC

**Data curation:** LQ XZ HZ FW QC

**Formal analysis:** LQ XZ HZ FW

**Funding acquisition:** CLQC

**Investigation:** LQ XZ HZ FW ML QC CL

**Resources:** CL

**Methodology:** LQ CL

**Supervision:** CL QC

**Project administration:** CL QC

**Visualization:** LQ XZ HZ FW CL

**Writing-original draft:** LQ ML CL QC

**Writing-review & editing:** LQ ML QC CL

## Expanded view figure legends

**Fig EV 1 -Elongator acts as an integral complex to regulate root growth.**

A Root length of different genotypes measured at 5 days after germination (DAG). At least 30 seedlings were measured for each genotype. Data shown are average and SD.

B Photograph of 5-day-old wild-type (WT) and *elp1* seedlings showing the short root of the *elp1* mutant.

C Root growth dynamics of the WT and *elp1*. Data shown are average and SD (*n* = 20).

D Root tips of the WT and *elp1* seedlings at 5 DAG, showing the significantly inhibited cell division and cell elongation in the *elp1* mutant. MZ, meristematic zone; EZ, elongation zone; DZ, differentiation zone.

E, F Statistics of the mature cortex cell length (E) and meristem cell number (F) at 5 DAG. Data shown are average and SD (*n* = 20). Asterisks indicate significant differences according to Student’s *t* test (** *P*< 0.01).

G, H *pCYCB1;1:GUS* expression in the WT (G) and *elp1*(H) at 5 DAG. Bars = 200 μm in (D) and 100 μm in (G) and (H).

**Fig EV 2 - Elongator acts independently of the PLT pathway.**

A Photograph of 5-DAG seedlings of the WT, *elp1*, *plt1-4 plt2-2*, and *elp1 plt1-4 plt2-2*.

B, E Root tips of the WT (B), *elp1* (C), *plt1-4 plt2-2* (D), and *elp1 plt1-4 plt2-2* (E) at 5 DAG. Vertical black lines indicate the root meristem of different plants extending from the QC to the transition zone.

F Quantification of the meristem cell number of the indicated plants. Data shown are average and SD (*n* = 20). Samples with different letters are significantly different at *P*< 0.01.

G, J Expression of *PLT1* ([G]and[H]) and *PLT2* ([I]and[J]) in 5-DAG WT ([G]and[I]) and *elp1* ([H]and[J]) revealed by whole-mount RNA *in situ* hybridization with a *PLT1* or *PLT2* antisense probe. Bars *=* 100 μm in (B) to (E) and 50 μm in (G) to (J).

**Fig EV 3- Subcellular localization of ELP3.**

APhotograph of 5-DAG seedlings showing that the *pELP3:ELP3-GFP* construct fully complemented the *elp3* mutant phenotype.

B Representative confocal image showing the subcellular localization of ELP3 in *p35S:GFP-ELP3* plants, noting more obvious nuclear localization in columella cells and epidermis cells at the elongation zone. Bar = 50 μm.

**Fig EV 4 - Antibody characterization and cell fractionation assay.**

A, B Validation of the anti-ELP1 (A) and anti-ELP3 (B) antibodies in Col-0 and *elp1* and *elp3*, respectively. Total proteins were extracted from 10-day-old Col-0 and *elp1* and *elp3* seedlings. Immunoblot was done with the anti-ELP1 and anti-ELP3 antibodies prepared in our laboratory. Ponceau S-stained membranes are shown as loading controls. Asterisk indicates the position of the specific band.

C Cell fractionation assay in *elp3;p35S:ELP3-myc* plants. Ten-day-old *elp3;p35S:ELP3-myc* seedlings were harvested for cell fractionation. Proteins from different fractions were immunoblotted with antibodies against *myc*, H3, and PEPC. H3 and PEPC were used as nucleus- and cytoplasm-specific marker proteins. T, total extracts; C, cytoplasmic fraction; N, nuclear fraction. The experiments were repeated three times with similar results.

**Fig EV 5 -Expression of *SPT4-1* and *SPT4-2* in *SPT4-RNAi* plants.**

A, B Expression of both *SPT4-1*(A) and *SPT4-2*(B) was significantly knocked down in *SPT4-RNAi* plants. Ten-day-old seedlings were harvested for RNA extraction and RT-qPCR analysis. The relative expression level was normalized to the reference gene *PP2AA3*. Error bars represent SD. Asterisks indicate significant differences according to Student’s *t* test (** *P*< 0.01).

**Appendix Table1 List of primers used in this study.**

